# Neural oscillatory dynamics in joint action: distinct contributions of entrainment and beta modulation to self–other integration

**DOI:** 10.1101/2025.04.23.650255

**Authors:** M. Rosso, B. Van Kerrebroeck, P. E. Keller, M. Leman, P. Maes, P. Vuust

## Abstract

Temporal coordination plays a pivotal role in human activities, enabling effective communication and collaboration. Yet, the neural mechanisms supporting this ability remain poorly understood. Recent research suggests that synchronized beta modulation across individual brains, as measured via EEG hyperscanning, may reflect a shared sensorimotor framework underpinning interpersonal synchronization. Building on the idea that self–other integration is a key mechanism in this process, behavioral studies have shown that adopting a partner’s first-person visual perspective (1P) enhances interpersonal coordination, as opposed to the natural second-person perspective (2P).

Here, we used the body-swap illusion in immersive virtual reality to directly manipulate embodied perspective and investigate its neural consequences. Specifically, we examined how this illusion modulates beta activity and neural entrainment during joint rhythmic action. Forty participants (N=40), randomly paired into twenty dyads, performed a joint finger-tapping task while wearing head-mounted displays for immersive virtual environments. In coupled conditions, the visual scene was manipulated so that participants viewed their partner’s hand from a 1P perspective, as if it belonged to their own body, or from a natural 2P perspective. In uncoupled control conditions, participants saw their own hand in 1P and 2P, without perceiving any information about their partner’s tapping.

EEG hyperscanning revealed that both neural entrainment (measured as the convergence of low-frequency oscillations) and beta modulation (changes in ∼20 Hz power linked to partner-generated movements) occurred in 1P and 2P visually coupled conditions. However, only beta modulation was selectively enhanced when participants experienced their partner’s hand from a 1P perspective. These findings suggest that while neural entrainment reflects a general mechanism for tracking a partner’s rhythmic behavior, beta modulation specifically supports the integration of the other’s effector into one’s bodily representation.

## Introduction

The ability to put oneself in another’s place is crucial for human cooperation and may be unique to our species. ^1^. This faculty, known in cognitive neuroscience as perspective-taking ^2^, holds more than just a metaphorical meaning. Evidence from a range of task domains suggests that perspective taking is an embodied process, whereby the brain integrates information from the other in relation to one’s own bodily state in order to infer their mental states and coordinate with their actions ^3–7^. Understanding how the brain facilitates this process is one of the primary quests of contemporary social neuroscience ^8^.

Perspective taking can be experimentally induced and investigated with the body-swap illusion ^9–11^. In this paradigm, the visual perspective of two participants is switched via head-mounted displays, such that one can perceive the other’s body from the other’s 1st person perspective (1P). This scenario differs from natural interpersonal interactions where individuals interact from a 2nd person perspective (2P), namely perceiving the other while maintaining one’s own point of view ^12^. This manipulation is particularly insightful because it is thought to trigger qualitatively different mechanisms of self-other integration ^13^, which is a critical component of human interactions ^14–17^. When the other’s body is perceived in 1P, due to the spatiotemporal congruency between the visual percept (the other) and the afferent proprioceptive signals (the self), the other’s body parts can be integrated into one’s own mental body schema ^11^. This process ties together perception and action relying on dynamic neural representations of the bodily-self, based on a sensorimotor recalibration of the body schema ^18,19^. While visual manipulations have been extensively used to investigate how the embodiment of external limbs comes with illusory sense of ownership and agency (for reviews, see ^20,21^), to our knowledge, no study to date has leveraged visual manipulations to uncover the neural dynamics underlying self-other integration and how these influence an ongoing social interaction.

Using the body-swap illusion, Rosso et al. ^13^ showed that inducing mutual perspective-taking between two individuals by switching their visual perspectives facilitates attraction towards a synchronized state, pointing to a dependency of temporal coordination from the integration of the other’s body state into the individual sensorimotor systems. The authors proposed that the assumption of a forward model for the other’s effector compels individuals to minimize the error between executed and observed action, which ultimately leads to an increase in behavioral synchronization ^22^. The aim of the present study is to go beyond behavioral evidence and understand the neural mechanisms by which the motoric information produced by another individual is integrated into the sensorimotor system during interpersonal synchronization.

In order to address this question, we applied two recently developed EEG analysis methods ^23–25^ to a hyperscanning dataset collected in the joint finger-tapping study of Rosso et al. ^13^. The task was carried out in a drifting metronomes paradigm, where each member of the dyads was assigned a metronome and instructed to synchronize their finger taps with it. While the two metronomes were set to start in phase, a small difference in their tempo resulted in repeated cycles of de-phasing. Despite the instruction to ignore the partner while maintaining synchronization with the assigned metronome, this setup has consistently resulted in unintentional synchronization among the partners across independent studies ^13,26,27^. Furthermore, the continuous de-phasing enabled the analysis of the time-varying aspect of the neural dynamics as the interaction unfolds, through phases of coupling and decoupling characterizing metastable behavior ^14,28–30^. This experimental design allowed us to quantify the emergence of the neural dynamics of interest as a result of visual coupling and, importantly, to compare them across 1P and 2P modes of interaction.

Based on a large corpus of literature, we identified two distinct oscillatory dynamics potentially underpinning low-level mechanisms of self-other integration. The first candidate mechanism is **neural entrainment**, consisting of the phase-alignment of a low-frequency brain oscillation with the behavior of the partner. Hyperscanning studies have traditionally relied on the construct of interbrain synchrony, which fails to capture its underlying mechanistic principles (for a recent critical review, see ^31^). In contrast, we explicitly modeled the convergence of individual oscillatory components toward a common frequency within the dyad, adhering to the precise definition of neural entrainment ^25,32,33^. Specifically, from each participant’s EEG signal, we extracted components attuned to the assigned metronome frequencies (1.667 Hz or 1.641 Hz) and computed the dynamic convergence of their instantaneous frequencies over time ^23,25,34^. To investigate these neural mechanisms, we manipulated Coupling (coupled and uncoupled) and Perspective (1P and 2P) in a within-subjects design, allowing us to systematically assess how visual coupling and embodied perspective-taking influence self-other integration.

For neural entrainment, we hypothesized that *A1)* visual coupling would result in significantly stronger frequency convergence of the individual oscillatory components, given that visual information is known to facilitate spontaneous interpersonal synchronization ^27,35–37^. Furthermore, we expected that *A2)* periods of frequency convergence would alternate with periods of frequency divergence, reflecting the alternating periods of coupled and decoupled behavior previously reported in the drifting metronomes paradigm. Finally, we hypothesized that *A3)* frequency convergence would be stronger in 1P as compared to 2P, consistent with the enhanced behavioral synchronization reported by Rosso et al. ^13^ and reflecting greater self-other integration.

The second mechanism of interest is **beta modulation**, consisting of changes in the power of a ∼20 Hz oscillatory component in response to rhythmic events. Notably, beta power modulation occurs both during self-generated movements ^38,39^ and in response to sensory inputs^40–43^, making it a candidate for regulating action timing in joint action ^24,44–46^. Operating at the interface of motor and sensory processes, beta modulation is thought to coordinate executed and perceived actions through a shared oscillatory mechanism ^24,47–49^. Beta modulation was operationalized as a form of brain-to-behavior coupling, which we quantified as variation in ∼20 Hz power as a function of the phase of the partner’s finger taps. We hypothesized that *B1)* visual coupling would result in significant beta modulation driven by the partner’s tapping cycles, and that *B2)* the modulation would be stronger in the 1P coupled condition as compared to the 2P coupled condition, reflecting enhanced integration of the partner’s movements into one’s own body schema due to embodied perspective-taking.

Together, these hypotheses address both a general principle of error minimization between self and other ^22^ through neural entrainment, as well as a specific sensorimotor integration via beta modulation. Support for these hypotheses would suggest that perspective-taking enhances self-other integration by modulating multiple levels of neural processing. Figure 1 provides a visual representation of the dynamics under investigation and our hypotheses, while Figure 2 illustrates the analysis pipeline leading to their computation.

**Figure 1.**
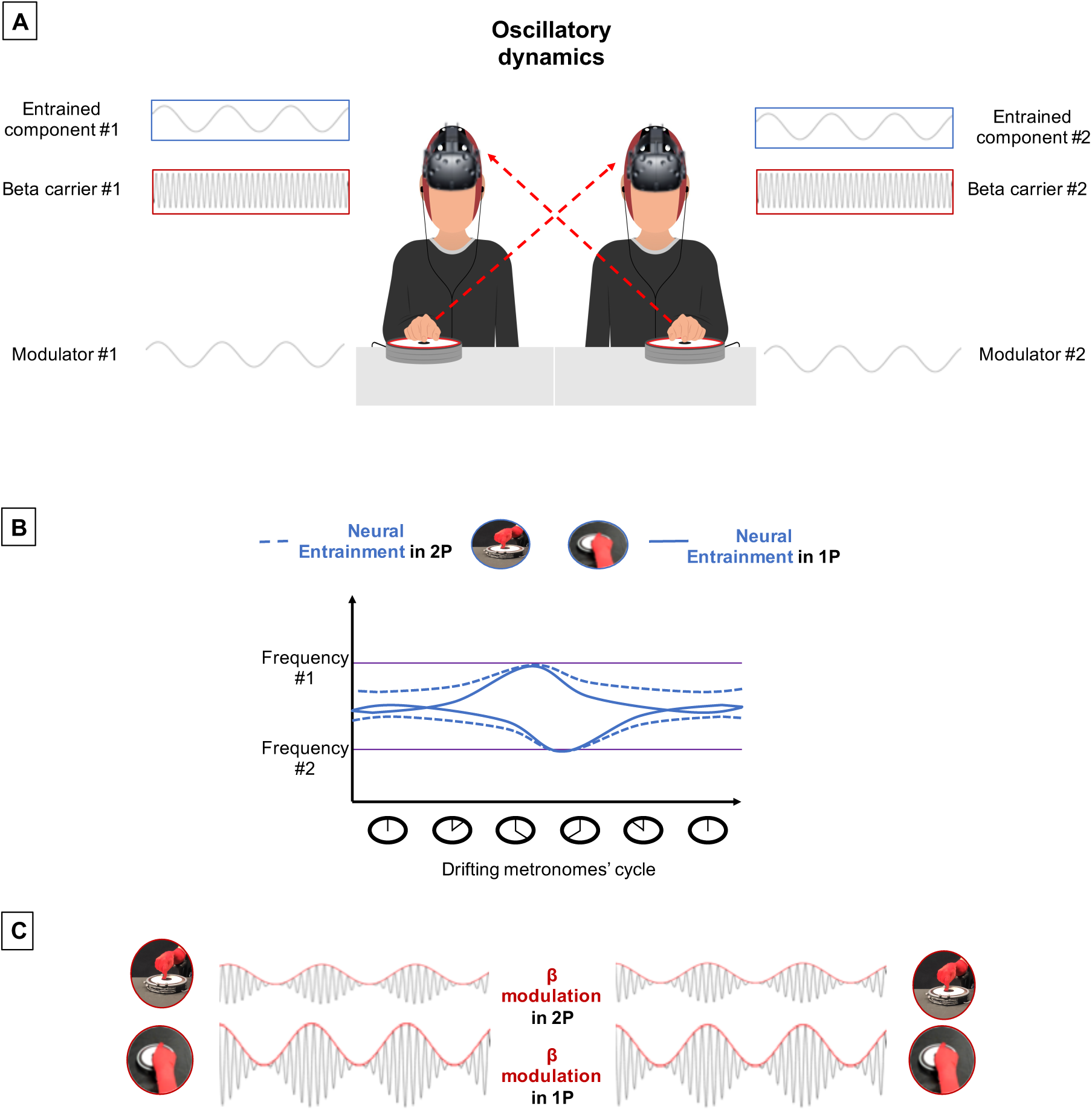
Schematic illustration of oscillatory neural dynamics underlying self–other integration during joint finger tapping. **(A) Oscillatory dynamics** under investigation are represented in the context of a joint finger-tapping task, where participants are visually coupled by observing each other’s hand through a head-mounted display (HMD). Participants observed their partner’s hand movements either from a first-person (1P) or a second-person (2P) perspective while synchronizing with temporally incongruent auditory metronomes (drifting metronomes). For each participant in the dyad, we aimed to separate the following oscillations (from top to bottom): a low-frequency component of the EEG signal attuned to the assigned metronome (entrained component), a high-frequency component centered at 20 Hz (beta carrier), and a low-frequency oscillation corresponding to the finger-tapping cycles (modulator). The body-swap manipulation affects how each participant processes the partner’s movements as if they were their own (1P), allowing to assess changes in the dynamics of interest compared to an ecological face-to-face interaction (2P). **(B) Neural entrainment** was quantified as frequency convergence between entrained components within dyads, across the drifting metronomes’ cycle. The two horizontal purple lines represent the ideal scenario in which each participant maintains their individual frequency based on the assigned metronome. The blue lines depict the expected alternating periods of frequency convergence and divergence across the cycle, modulated by visual coupling and perspective (dashed and solid lines represent 2P and 1P, respectively). **(C) Beta modulation** reflects periodic fluctuations in the power of ∼20 Hz oscillations driven by the partner’s tapping. Modulation strength is expected to be higher in the 1P perspective, based on enhanced integration of the other’s effector into the self’s body schema as compared to an ecological 2P perspective. The curves are intended for illustrative purposes only.

**Figure 2.**
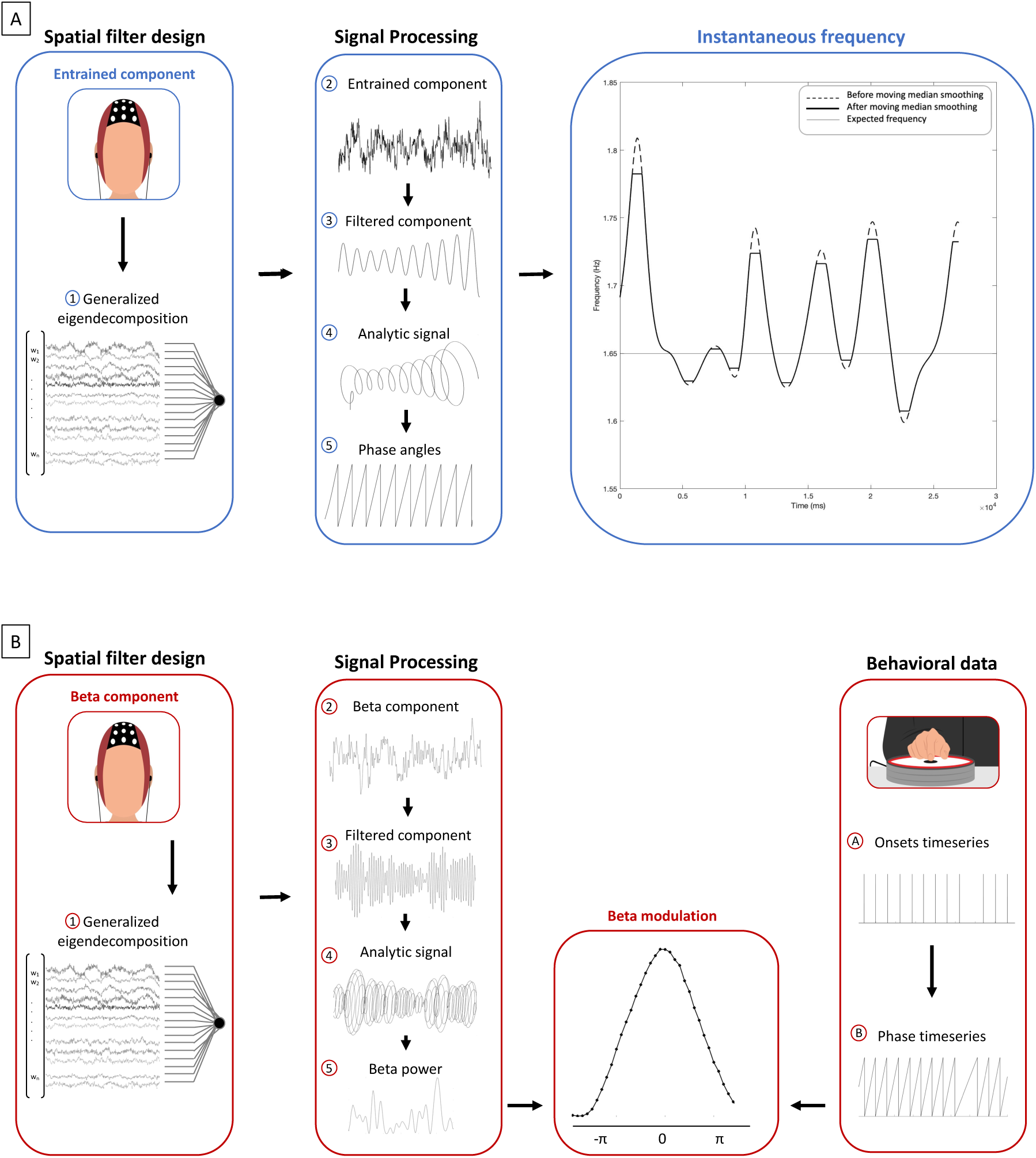
Signal processing pipelines for extracting oscillatory components from EEG hyperscanning data. **(A) Neural entrainment pipeline.** A spatial filter was computed using generalized eigendecomposition (GED) (1) to extract the EEG component most attuned to the metronome’s frequency. The extracted entrained component (2) was filtered (3), and its analytic signal (4) was computed via Hilbert transform to extract instantaneous phase angles (5). Based on the rate of change of the phase time series, **instantaneous frequency** was derived and smoothed with a moving median filter. Carrying out this pipeline for each participant in the dyad allowed estimation of frequency changes over time, and their convergence across the metronomes’ cycle. **(B) Beta modulation pipeline.** A spatial filter was computed using generalized eigendecomposition (GED) (1) to extract an EEG component that best separates narrow-band beta activity (∼20 Hz) from broadband activity. The extracted beta component (2) was filtered (3), and its analytic signal (4) was computed via Hilbert transform to extract beta power time series (5). In parallel, behavioral onset time series (A) from the partner’s finger taps were converted into phase time series (B). Beta **power** was then binned as a function of tapping phase, and a sinusoidal function was fit to quantify the strength of beta modulation driven by the partner’s tapping cycles.

## Results

We found that both neural entrainment and beta modulation were elicited when partners performed the finger-tapping task in visually coupled conditions. However, each mechanism responded selectively to our manipulation of visual perspective, suggesting they play distinct roles in the integration of motoric information produced by the partner during interpersonal synchronization.

### Neural entrainment

Neural entrainment was quantified as the convergence of instantaneous frequencies between the partners’ entrained EEG components over time. We found a significant 2-way interaction effect between Coupling and the quadratic term of Time (Estimate = 2.446, SE = 0.005, p < 0.01), indicating that frequency convergence in coupled conditions was modulated as a function of the drifting metronomes’ cycle. As shown in Figure 3, the oscillations maximally converged toward a shared frequency around the in-phase attractor point, and diverged toward the respective metronome’s frequencies after the antiphase point. This parabolic modulation enabled us to model the time-varying component of convergence using polynomial fitting ^50^ (for more details on the statistical model, see the Methods section). Furthermore, we found a significant 2-way interaction between Coupling and the linear term of Time, capturing the symmetry between the rates of divergence and convergence (Estimate = -0.024, SE = 0.006, p < 0.01). As shown in Figure 3, frequencies reached maximum divergence after the anti-phase midpoint in the coupled 2P condition, followed by a steeper attraction as they re-entered in phase at the other end of the cycle. This pattern closely resembles the asymmetric behavioral coordination dynamics consistently reported in independent studies using the drifting metronomes paradigm ^13,26,27^. Specifically, the in-phase attractor exerted a longer-lasting influence on neural convergence as participants attempted to de-phase away from its region. Against our initial hypothesis, we did not find any significant interaction effect between Perspective and the quadratic component of the modulation, indicating that the depth of frequency convergence was invariant between 1P and 2P coupling. However, the significant 3- way interaction between Coupling, Perspective, and the linear term of Time (Estimate = 2.248, SE = 0.009, p = 0.014) shows that visual perspective influenced the asymmetry of the dynamic described above. When participants were coupled in 1P, the rates of divergence from and convergence toward the in-phase point became significantly more symmetric. Table 1 shows the fixed-effects parameter estimates, their standard errors, and associated p-values. Uncoupled (factor Coupling) and 2P (factor Perspective) were taken as 0-levels for statistical contrasts.

**Figure 3.**
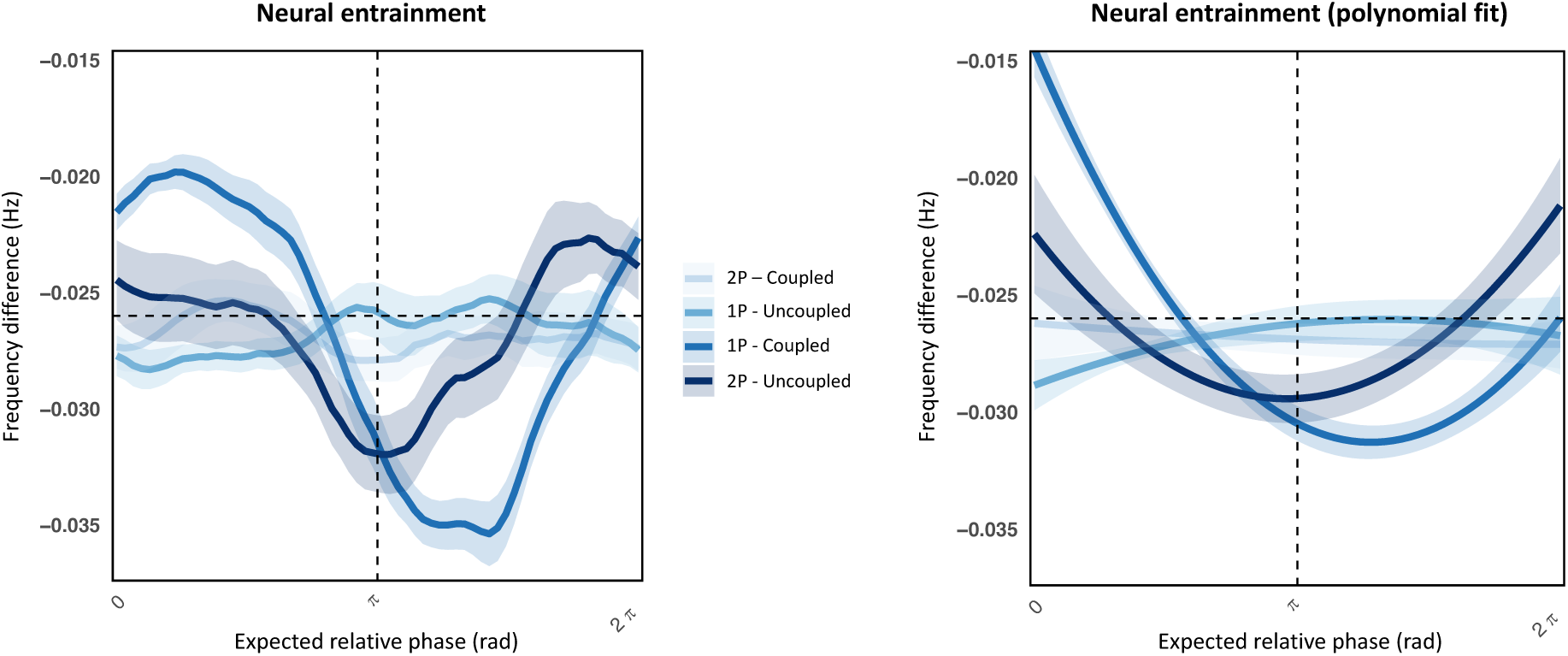
Neural entrainment as frequency convergence. The plot on the left shows neural entrainment quantified as the difference in instantaneous frequency (Hz; y-axis) between the entrained components of each partner’s EEG signal, computed over the drifting metronomes’ cycle (x-axis, expressed as expected relative phase in radians from 0 to 2π). Continuous lines indicate the average frequency difference computed from all drifting metronomes’ cycles and from all participants (N = 36); ribbons indicate the standard error of the mean (SEM) across participants. Values closer to zero indicate stronger convergence between partners’ neural signals. Across all visually coupled conditions, oscillatory components converged toward a shared frequency around the in-phase attractor point and diverged after the anti-phase midpoint. As expected, both 2P and 1P Uncoupled conditions fluctuated around -0.026 Hz, which is the expected difference based on the difference in the metronomes’ frequencies. Instead, both 2P and 1P Coupled conditions showed a clear parabolic pattern of convergence, with a significant quadratic modulation. Against our predictions, the overall depth of convergence remained unaffected by perspective, as quantified by the non-significant interaction between the quadratic term of the model and Coupling. However, 2P coupling exhibited more asymmetric convergence-divergence slopes, while the 1P perspective rebalanced this asymmetry, leading to more symmetric convergence dynamics. The plot on the right represents the quadratic polynomial fit to the same curves, highlighting the asymmetry in convergence across 1P and 2P Coupled conditions.

**Table 1.** Neural entrainment results. Results from a linear mixed-effects model examining neural entrainment as a function of drifting metronomes’ cycles (ot1 and ot2, polynomial terms of Time), visual perspective (1P vs. 2P), and movement coupling (Coupled vs. Uncoupled). The model was fitted to the frequency difference over the drifting metronomes’ cycle, and includes fixed main effects and interactions. Significant interaction effects between factors and polynomial terms indicate modulation of neural phase alignment by sensorimotor perspective and interpersonal coupling. Significance thresholds: p < .05 *, p < .01 **, p < .001 ***.

| Term | Estimate | Std. Error | t value | p value | Significance |
| --- | --- | --- | --- | --- | --- |
| (Intercept) | -0.027 | 0.000 | -50.534 | 0.000 | *** |
| ot1 | -0.002 | 0.004 | -0.445 | 0.656 |  |
| ot2 | 0.001 | 0.004 | 0.139 | 0.895 |  |
| Perspective | 0.000 | 0.001 | 0.062 | 0.950 |  |
| Coupling | -0.001 | 0.001 | -0.196 | 0.844 |  |
| ot1:Perspective | 0.007 | 0.006 | 1.122 | 0.262 |  |
| ot2:Perspective | -0.004 | 0.005 | -0.890 | 0.373 |  |
| ot1:Coupling | -0.025 | 0.006 | -3.899 | 0.000 | *** |
| ot2:Coupling | 0.024 | 0.005 | 4.818 | 0.000 | *** |
| Perspective:Coupling | 0.001 | 0.001 | 0.121 | 0.903 |  |
| ot1:Perspective:Coupling | 0.022 | 0.009 | 2.467 | 0.014 | * |
| ot2:Perspective:Coupling | -0.002 | 0.007 | -0.319 | 0.749 |  |

### Beta modulation

Beta modulation was quantified as the amplitude of the beta power distribution over the partner’s finger-tapping cycles. We found a significant 2-way interaction between Coupling and Perspective (Estimate = 0.371, SE = 0.178, p = 0.037), indicating that coupling with the partner in 1P resulted in significantly stronger beta modulation as compared to 2P. This effect extends the findings reported by Rosso et al. ^24^, showing that a larger allocation of neural resources is devoted to tracking the partner’s effector when it is perceived from a perspective that facilitates integration into one’s own body schema. The results and interactions in the model are represented in Figure 4. Table 2 contains the full list of contrasts performed across the model’s posterior distributions. Uncoupled (factor Coupling) and 2P (factor Perspective) were taken as 0-levels for statistical contrasts.

**Figure 4.**
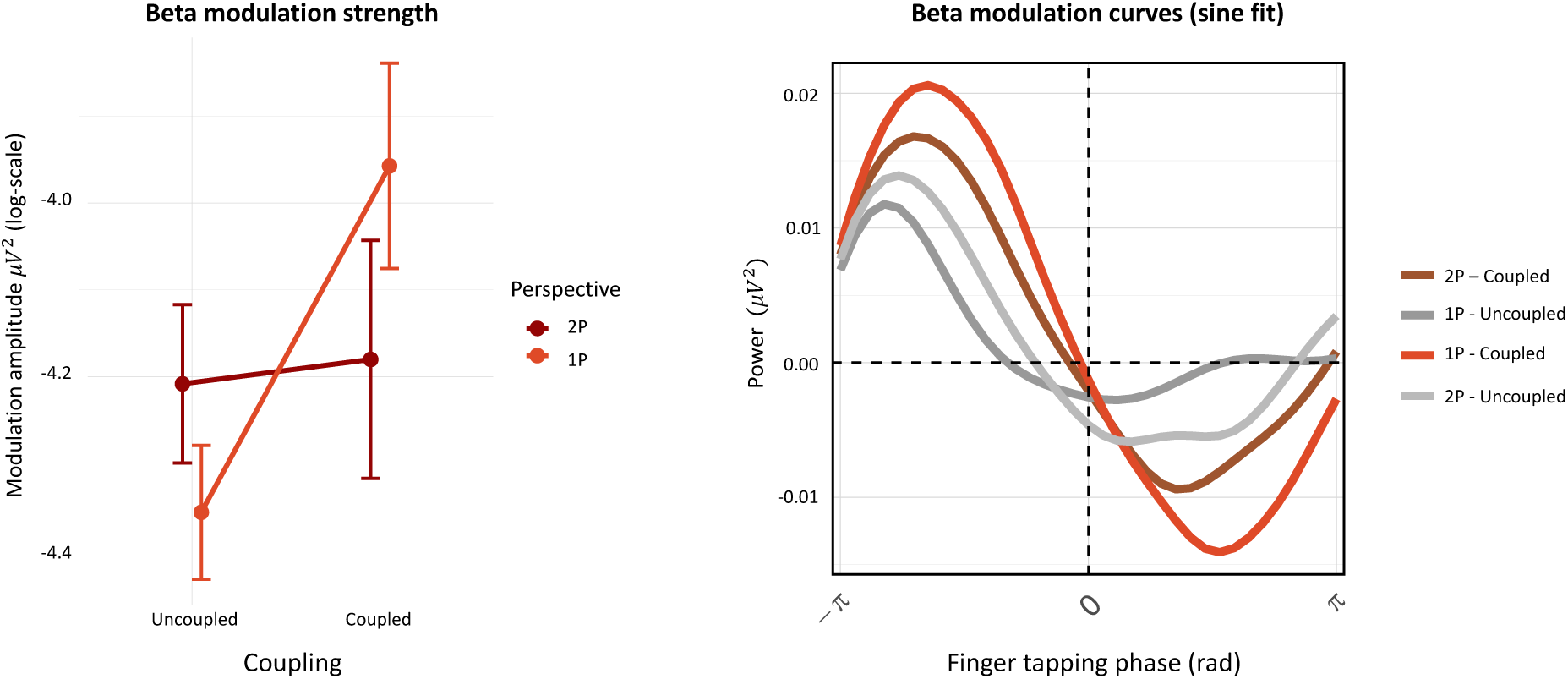
Beta modulation as a function of the partner’s movement cycles. The plot on the left shows the average **beta modulation strength** computed across 36 participants (N = 36) in each experimental condition. Mean amplitude of beta-band modulation (log-transformed; y-axis) is shown in response to the partner’s tapping cycle, plotted across levels of Coupling and color-coded by Perspective (2P vs 1P). Beta modulation was quantified by fitting a sine function to the time-varying beta power aligned to the partner’s tapping phase. A significant interaction was observed between Coupling and Perspective, driven by a larger increase in beta modulation in the 1P Coupled condition compared to 2P. This suggests enhanced beta dynamics when participants viewed the partner’s movements from a first-person perspective while visually coupled. Error bars represent standard errors of the mean (SEM). The plot on the right shows the corresponding group-averaged **beta modulation curves**, aligned to the partner’s tapping phase (x-axis, in radians), with a fitted sine wave for each condition. The curves reveal the temporal structure of beta power modulation, peaking and troughing in alignment with the partner’s movement cycle. The 1P Coupled condition exhibits the clearest oscillatory profile, consistent with increased modulation strength, while the other conditions show reduced or more diffuse modulation patterns.

**Table 2.** Beta modulation results. Results from a linear mixed-effects model predicting the strength of beta-band modulation (∼20 Hz) by behavioral tapping phase. The model includes fixed effects of visual perspective, movement coupling, and their interaction. Significant beta modulation in the Coupled × 1P condition suggests enhanced integration of the partner’s tapping cycles into the self’s motor representation when visual perspective is first-person and sensorimotor coupling is present. Significance thresholds: p < .05 *, p < .01 **, p < .001 ***.

| Term | Estimate | Std. Error | t value | p value | Significance |
| --- | --- | --- | --- | --- | --- |
| (Intercept) | -4.209 | 0.107 | -39.333 | 0.000 | *** |
| CouplingCoupled | 0.028 | 0.127 | 0.223 | 0.824 |  |
| Perspective1P | -0.149 | 0.127 | -1.172 | 0.242 |  |
| CouplingCoupled:Perspective1P | 0.372 | 0.179 | 2.077 | 0.038 | * |

To summarize, neural entrainment as a result of visual coupling manifested as the frequency convergence of individual oscillatory components initially attuned to the assigned metronomes. Across the drifting metronomes’ cycle, this dynamic was susceptible to the influence of attractor points, alternating between phases of convergence and divergence. Convergence strength was invariant to visual perspective, while the convergence and divergence slopes were more similar in 1P coupling. Beta modulation strength, on the other hand, increased significantly when participants were coupled in 1P compared to 2P. The differences in the susceptibility of neural entrainment and beta modulation to visual perspective point toward a functional specialization of the two mechanisms underlying interpersonal synchronization.

## Discussion

With the present work, we showed that distinct oscillatory neural dynamics underpin distinct functions in interpersonal synchronization, integrating different dimensions of the motoric information produced by a human partner. While both neural entrainment and beta modulation were elicited in all conditions of visual coupling, beta modulation was selectively enhanced when participants perceived their partner’s hand from a 1st person perspective (1P), as if it belonged to their own body. Our results suggest that while neural entrainment reflects a general mechanism for tracking a partner’s rhythmic behavior, beta modulation selectively supports the integration of the other’s effector into one’s bodily representation.

By examining these mechanisms under the body-swap illusion ^9,10^, we have shown that beta modulation, in particular, is enhanced under conditions that induce a recalibration of the sensorimotor system, such as perceiving the partner’s hand from a 1st person perspective (1P) (see Figure 4). This enhanced modulation suggests that beta rhythms are not merely a generalized mechanism for tracking rhythmic stimuli ^40,41,52,53^ and perceived actions ^24,42,45,46,49^. Instead, they might play a crucial role in dynamically integrating spatiotemporal information about the current body state to update one’s body schema ^18,19,54^. The alignment of visual and proprioceptive cues within the 1P condition creates the minimal and sufficient conditions needed for incorporating an external body part into a coherent sense of self-location and body ownership ^55–57^. Importantly, this integration is strengthened by visuo-motor synchronization, namely when the perceived effector moves in congruence with the observer’s motor commands and sensory predictions ^58–60^. We propose that rhythmic beta modulation facilitates this integration by aligning with the phase of the partner’s movements.

Beta modulation has been shown to differentiate contributions by the self and the other to joint action within the motor system based on different degrees of beta power suppression ^49^, and has been proposed as a shared oscillatory mechanism for pacing one’s own movements as well as tracking those of others ^24^. Noteworthy, the increment in modulation in 1P suggests that such differentiation no longer holds in a condition where the partner’s actions are integrated into the self’s body-schema. The concept of self-other integration is paramount to interpersonal coordination, as integrating motoric information from another person’s body into the perceiver’s motor program enables individuals to adjust their actions in response to the other’s movements ^14,15,22,61,62^. As a prominent endogenous rhythm in the motor cortex and supplementary motor area ^63,64^, beta oscillations are well-positioned to support this integration due to their role in sustaining motor execution and control through corticomuscular coupling^65^. Beta rhythms uniquely coordinate both efferent motor commands along the corticospinal pathway and afferent signals that inform the brain about the body’s current state, enabling continuous recalibration and stabilization ^66–69^. This dual function of beta oscillations, encoding outgoing and tracking incoming signals, makes them ideally suited to handle the dynamic requirements of incorporating another’s movements into one’s own motoric framework. Novembre et al. ^44^ previously reported that inducing 20 Hz inter-brain synchrony between two individuals’ motor cortices aligned sensorimotor processing within the dyad. The alignment of excitability phases of beta oscillations between the two brains influenced the timing of joint action initiation and improved the likelihood of achieving interpersonal movement synchrony. This finding suggests that beta oscillations play a causal role in coordinating motor actions between interacting individuals ^70^. While the authors specifically focused on phase-alignment, they put forward that an envelope with a 20 Hz carrier modulated by a movement frequency, as we computed in our study, would account for the contextual timing of the dyadic interaction ^44^. According to this view, simultaneous movement would be promoted by alternating windows of motor excitability and inhibition coupled to the partner’s tapping ^71,72^. In line with this evidence, we propose that sensorimotor integration aligns efferent motor commands to the timing of the other’s effector, compelling coupled individuals toward a synchronized state and thereby minimizing the mismatch between executed and observed actions ^22^.

Unlike beta modulation, which facilitates self-other integration, our findings indicate that neural entrainment supports synchronization with external stimuli without incorporating them into the self’s motor schema. This distinction was highlighted by the body-swap illusion, where the frequency convergence of entrained EEG components remains unaffected, suggesting that neural entrainment operates independently from sensorimotor recalibration. Instead, it tracks the partner’s effector as a distal stimulus rather than integrating it into the self-schema. By aligning the frequency of internal oscillations to the other’s rhythmic movement, neural entrainment enables sensorimotor synchronization ^25^ without extending into representational processes.

In our study, the entrained oscillations under analysis fall within the delta range (0.5–4 Hz), an intrinsic oscillatory rhythm prominent in the motor system alongside beta rhythms ^63,73^, which aligns with the natural rhythm of human motion as constrained by bodily limitations ^74^. Since observing another person moving naturally overlaps with the perceiver’s endogenous delta rhythms in motor areas, the proximity in frequencies renders human rhythmic motion a unique class of social rhythmic stimulus which provides a natural affordance for the motor system to entrain ^75^. In order to set the initial conditions for individual brains to converge toward a shared frequency, our drifting metronomes paradigm introduced a frequency gap between participants’ assigned metronomes. By extending recent advances in multivariate EEG analysis ^23,25^ to hyperscanning recordings, we operationalized neural entrainment in the interpersonal domain following its fundamental definition ^32,76,77^. Specifically, we separated each participant’s oscillatory component most attuned to their metronome and tracked its instantaneous frequency changes throughout the task, to quantify convergence and divergence dynamics as the drifting metronomes’ cycle unfolded.

As shown in Figure 3, in both 2P and 1P visually coupled conditions, these oscillatory neural components diverged from the assigned metronome’s frequency, gravitating instead toward the partner’s frequency as the dyad approached the in-phase attractor in the drifting metronomes’ cycle. Around the anti-phase midpoint, partners de-coupled, realigning their oscillatory component with their own metronome. This cyclical pattern of convergence and divergence, shaped by attractor points in the drifting metronomes’ cycle, mirrors the same attractor landscape underlying behavioral synchronization ^13,26,27^, suggesting that neural entrainment is a core mechanism in this process. While the overall extent of frequency convergence was consistent across visual perspectives, the balance between convergence and divergence rates varied. Specifically, coupling in 2P conditions produced a slower divergence followed by a steeper convergence, whereas 1P conditions balanced these dynamics, eliminating the directional effect seen in 2P (or ‘hysteresis’ ^78^). This nuanced dynamic does not alter our conclusions on the extent of the convergence, but it suggests subtle differences based on visual perspective that could be further explored through computational modeling of dwelling times, hysteresis, and memory effects in the system, to better clarify the underlying mechanisms. Importantly, a similar rebalancing was empirically observed at the behavioral level ^13^, reinforcing the link between neural entrainment and behavioral synchronization.

In line with prior findings ^73,79,80^ and with the dynamic attending theory ^81,82^, we propose that cortical delta dynamics ^83–85^, particularly their phase ^86–89^, regulate attentional focus on competing rhythmic sensory inputs, namely, the assigned metronome versus the partner’s movements. Translated to the interpersonal domain, participants appear to track each other’s movements and guide their own behavior toward synchronized states through frequency convergence. This framework implies that dynamic allocation of attentional resources to competing inputs mediates transitions between cooperation and competition processes ^26,28^, otherwise described as metastable behavior within the dyad ^14,29^. While our experiment cannot establish causality between oscillatory brain mechanisms and individual or dyadic behavior, the motor origin of delta oscillations has led authors to argue that active behavioral engagement drives cortical phase alignment, thereby enhancing sensory prediction ^73,84,90^. In line with this argument, our results suggest that neural entrainment may facilitate prediction error minimization by reducing the temporal mismatch between executed and observed actions ^22^. To conclude, our findings highlight that distinct modulation mechanisms integrate distinct action components perceived from another human in order to support interpersonal synchronization. These mechanisms are frequency modulation for delta (neural entrainment) and power modulation for beta (beta modulation) rhythms. Each frequency range plays a specific role in mediating temporal coordination between individuals, with delta oscillations tracking general rhythmic alignment and beta selectively recalibrating the sensorimotor representation of the other’s effector. This distinction underscores how temporal synchronization between humans relies on qualitatively different neural processes that work together to achieve interpersonal coordination.

### Limitations and future directions

Despite the novel contributions of this study, some limitations must be acknowledged. In particular, our ability to infer the precise brain networks underlying the observed oscillatory dynamics is constrained by the limited spatial resolution of EEG. Consequently, our discussion focused on temporal dynamics without speculating on the spatial extent of the neural networks involved. Future studies should take this limitation into account, and apply different data acquisition methods and analytical approaches to overcome it. We point out that the recent development of *network estimation via source separation* (*NESS*) provides a promising path forward to integrate the temporal and spatial levels of analysis, by separating the sources of oscillatory activity in anatomical source space. Now adapted for frequency-resolved (FREQ-NESS ^64,91^) and broad-band (BROAD-NESS ^92^) analysis of whole-brain voxel data reconstructed from magnetoencephalography (MEG), this framework could enable future studies to capture the spatiotemporal characteristics of brain networks engaged in self–other integration during social interactions.

In addition, an alternative interpretation of our beta modulation findings relates to the perception of sensorimotor mismatches. Rather than reflecting the integration of an external effector per se, enhanced beta modulation in the 1P condition could stem from increased sensitivity to mismatches between action and perception, which are expected to be strongest in 1P visual coupling. These mismatches may disrupt phase-aligned synchronous activity or alter phase-resetting dynamics in neural populations. While our experimental design focused on dyadic interaction, future studies could test this alternative hypothesis in single-participant setups, by introducing dynamic delays in the visual feedback of one’s own movements to isolate the influence of perceived sensorimotor discrepancies from the integration of external body parts.

Importantly, while our experimental manipulations effectively elicited the oscillatory neural dynamics of interest, the findings remain correlational and therefore do not allow strong conclusions regarding their causal influence on dyadic behavior. Establishing causal links between specific neural mechanisms and behavioral outcomes will require intervention-based approaches such as neurostimulation ^44,70^ or computational modelling.

Finally, from an applied perspective, by selectively enhancing beta modulation over neural entrainment using the body-swap illusion, we demonstrated how technology-mediated interactions can target specific neural processes. This empirical evidence lays the groundwork for developing interventions tailored to individual needs in fields such as physical therapy, neurorehabilitation, sports, and music training.

### Conclusion

Our work builds on a line of behavioral research in interpersonal synchronization, which has shown that interactions can be shaped by manipulating the flow of information between individuals ^13,26,27,35–37^. By extending recent developments in multivariate EEG analysis ^23–25,93,94^ to a hyperscanning context, we delineated the distinct functional roles of neural entrainment and beta modulation in interpersonal synchronization.

To conclude, we underscore that rhythmic social interactions naturally induce diverse forms of neuromodulation by engaging mechanisms such as neural entrainment and beta modulation. These findings contribute to understanding the neural mechanisms that support interpersonal coordination and highlight how they can be modulated by changes in perceptual coupling.

## Methods

### Participants

Forty (N = 40) right-handed human participants, recruited through university mailing lists and local Facebook groups took part in the study (28 females, 12 males; mean age = 31.42 years, standard deviation = 7.49 years). Individuals were divided into two gender-matched groups and randomly paired in twenty dyads (N = 20), in order to control for gender bias in the interaction. Two dyads were excluded from the analysis: one because of failure to comply with the instructions, one because a participant requested to perform the behavioral task without undergoing EEG recording. None of the participants had history of neurological, psychiatric or major medical disorders, nor were professional musicians. All participants declared they did not know the assigned partner before the experiment took place. The study was approved by the Ethics Committee of Ghent University (Faculty of Arts and Philosophy) and informed written consent was obtained from each participant, who received a 20 € coupon as compensation for their participation.

### Experimental task

The behavioral task consisted of the drifting metronome paradigm for dyadic entrainment, a joint-finger tapping task described in detail in ^13,26,27^. The two partners in the dyad were sitting at the same table, facing one another, and were instructed to tap their right index finger on a circular pad placed in front of them, in sync with an auditory metronome. Each partner was cued with a metronome set at a slightly different tempo (100 BPM and 98.5 BPM, or 1.667 Hz and 1.641 Hz), resulting in a pattern of linear de-phasing. Specifically, the metronomes’ relative phase gradually increased from 0 to 𝜋 radians, and then decreased from 𝜋 to 0 radians over the course of 10 consecutive cycles, for a total duration of 390 seconds. Participants were instructed to ignore any visual input and keep tapping along with the assigned metronome. During the task, participants were equipped with HTC Vive Pro 2 head-mounted displays for immersive virtual reality environments, which we used to stream video recordings of their own (Uncoupled conditions) or of the partner’s hand (Coupled conditions) in real time, from a 1st person (1P conditions) or 2nd person (2P conditions) perspective. The combination of these factors resulted in the 2 x 2 experimental design (Coupling x Perspective). We refer to ^13^ for more technical details of the system setup, validation of the visual perspective manipulation in VR, technical details of behavioral data capture and stimuli presentation, and additional data collected from the participants which are not reported in the present work.

### Data acquisition

Each participant was provided with a circular pad containing a strain gauge pressure sensor, used to detect tapping onsets with a 1 ms resolution. For each dyad, two pads were connected together to a Teensy 3.2 microcontroller, which worked as a serial/MIDI hub to log finger-tapping onsets and communicate with the experimental PC and EEG system. Simultaneous EEG recordings were performed on both partners within each dyad during the entire experiment. At the beginning of each metronomes’ cycle, a TTL trigger was sent from the Teensy microcontroller to the EEG amplifiers via BNC connection, to synchronize behavioral and neural timeseries. Each participant was equipped with a 64-channels waveguard TM original EEG headset (10-10 system, with Ag/AgCl electrodes). Two ANT-Neuro eego TM mylab systems were connected with a trigger adapter cable provided by the manufacturer to synchronize the recordings. Recordings were performed at a sampling rate of 1 kHz. Each pair of headsets shared a common ground, while “CPz_partner1” was used as common reference electrode for both of the partners. Impedances were monitored in the eego TM software environment and kept below 20 k Ω.

### EEG pre-processing

The EEG pre-processing pipeline was written integrating functions from the Fieldtrip toolbox ^95^ for Matlab (MathWorks Inc, USA). For every dyad, data were acquired as a 128-channels recording and split offline in 2 x 64-channels subsets, individually re-referenced to the electrodes ‘CPz_partner1’ and ‘CPz_partner2’, respectively. Following the rejection of bad channels, identified based on visual inspection of the raw timeseries and on the distribution of variance across channels, we re-referenced the two recordings to the average of the respective 64 electrodes. This procedure prevented extreme noise from the removed bad channels to leak into the common average. On average, 4.9 bad channels (SD = 2.9) were removed per participant per experimental condition. A sixth-order Butterworth high-pass filter with 1 Hz cut-off was applied to the raw recordings to remove slow drifts. This conservative threshold was taken given the long duration of the recordings and the motor task involved. As shown in ^25^, these parameters of the high-pass filter do not influence the oscillatory dynamics relevant to neural entrainment. A low-pass sixth-order Butter- worth filter with 40 Hz cut-off was applied to remove high-frequency muscular activity. A fourth-order notch filter centered at 50 Hz was applied to remove power-line noise up to the 3rd harmonic.

Subsequently, independent component analysis (ICA) ^96–98^ as implemented in the ‘runica’ Fieldtrip algorithm was conducted. Stereotyped artifacts were identified by means of visual inspection of the components’ scalp topographies and activation timeseries. The reference ‘CPz’ and the bad channels’ timeseries were excluded from the input data matrix. Under optimal conditions, removal would have been limited to the components exhibiting the stereotypical frontal distribution generated by blinks and lateral eye movements, or bilateral temporo-mastoidal distribution with periodic peaks in the activation timeseries attributable to heart beats. However, as compared to our previous report on the pre-processing of hyperscanning data in the same experimental paradigm ^24^, we opted for a more stringent approach and removed abnormal independent components characterized by sustained high-frequency noise or recurrent transients in activation, and typically centro-frontal clusters. We point out that some of this artifactual components might be caused by interference of the setup with the HMDs, and call for extra caution when cleaning EEG data collected under these conditions. However, we can safely conclude that eventual residual contributions of these artifacts were kept to a minimum, because centro-frontal clusters were excluded from the macro selection of regions of interest (ROI), while we reported significant differences across conditions which cannot be explained by residual noise. On average, 6.3 artifactual components (SD = 3.6) were identified and removed per participant per experimental condition. A visual comparison of the dataset before and after IC removal was carried out to assess the quality of the removal. Special attention was given to the frontal clusters of electrodes maximally contaminated by eye-related artifacts. Rejected bad channels were reconstructed after artifact removal, by computing the average activity from neighboring electrodes indicated by the template provided by ANT-Neuro for 64-channel waveguard TM original caps. No segmentation in epochs was performed during this pre-processing phase, so that every experimental condition was treated as a continuous experimental run.

### Generalized eigendecomposition

Generalized eigendecomposition (GED) ^94^ was used to design a spatial filter in order to separate narrow-band activity in the frequency ranges of interest from the background broad- band activity ^99^, while reducing the dimensionality of the multivariate EEG dataset to a single timeseries (details presented below). These extracted component activation timeseries (i.e., the estimation of the underlying source’s activation) were used as input for the hyperscanning analyses presented in the next paragraph. Replicating the analysis pipelines adopted in our previous work, we extracted the entrained component maximally attuned to the stimulation frequency (1.667 Hz or 1.641 Hz, depending on the assigned metronome) and a narrow-band beta component (20 Hz) from the individual EEG recordings. For the entrained components, the filter was designed in the frequency domain as a Gaussian function with central frequency of 1.667 Hz or 1.641 Hz (depending on the assigned metronome) and full width at half the maximum of 0.3Hz ^23,25^. For the beta component, the filter was designed as a plateau-shaped finite impulse response (FIR) filter centered at 20 Hz and covering a frequency range of 18-22 Hz (slope = 15%) ^24^. For both components, the broad-band EEG data were filtered via element-wise multiplication between the spectrum of the broadband signal and the spectrum of the narrow-band filter kernel, before transforming the resulting spectrum back in the time domain with inverse Fourier transform.

The EEG datasets were spatially filtered by weighting the channels timeseries. The set of vectors W (weights) was calculated by solving the following equation:

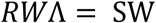

where S is the covariance matrix of the narrow-band signal; R is the reference covariance matrix of the broad-band signal; Λ is the set of eigenvalues. GED identified the set of eigenvectors W that best separate the signal (‘S’) covariance from the reference (‘R’) covariance matrix. The S covariance matrix was computed from the narrow-band signal, the R covariance matrix was here computed from the broad-band multivariate signal.

The eigenvector (w) associated to the largest eigenvalue was taken as a spatial filter, transposed and multiplied by the broadband data matrix to reconstruct the activation time series (y) of our target components:

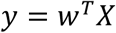

The corresponding spatial projection of the components over the scalp (a) were computed as follows:

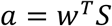

Covariance matrices were computed within 600 ms time windows starting at the finger-taps onsets, and grand-average S and R covariance matrices were computed. Matrices whose z-normalized Euclidean distance from the grand-average exceeded the 2.23 z-scores (corresponding to a probability of 0.01) were removed, and the grand-average S and R were recalculated based on the remaining covariance matrices. To improve the numerical stability of the GED performance in case of rank deficiency, regularization was applied to the R covariance matrix by adding 1% of its average eigenvalues as a constant to its diagonal.

Besides optimizing the signal-to-noise ratio between narrow- and broad-band neural activity, GED allows to avoid channel selection bias. Aiming to replicate our previous work, the spatial filter was optimized based on a macro-selection of regions of interest (ROIs), consisting of a large cluster of the 37 channels located behind the frontocentral ‘FC’ line - mastoids excluded. The S and R covariance matrices were computed from this cluster of ROIs. The rationale was to provide full coverage of centroparietal, temporal and occipital regions where the dynamic of interest were shown to be maximally expressed, while minimizing noise from less relevant sensors ^23,24^.

A quality assessment of the GED source separation is provided in Figure 5.

**Figure 5.**
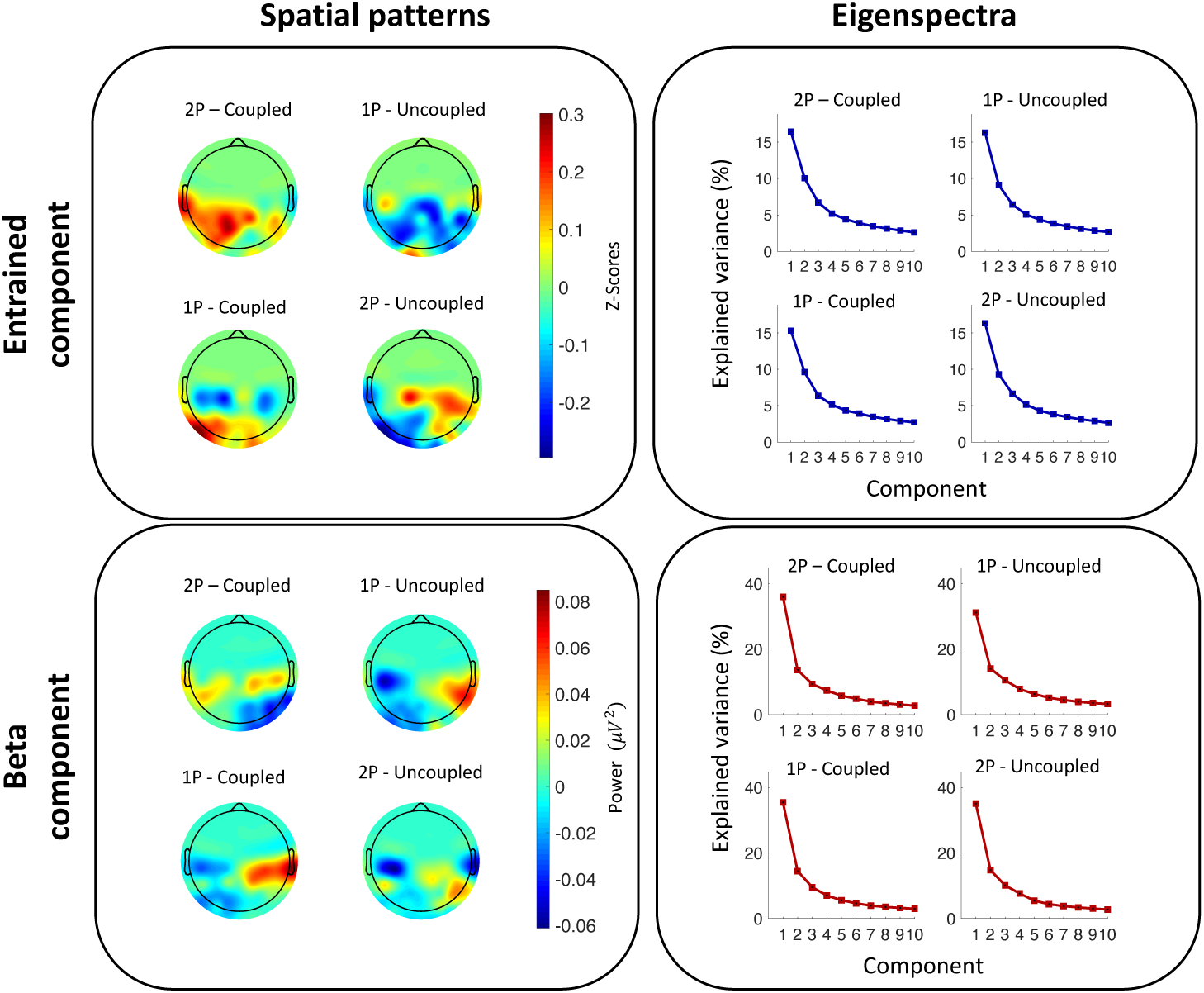
GED assessment. For both the entrained component and the beta component, spatial activation patterns and eigenspectra are shown across the four experimental conditions (2P–Coupled, 1P–Uncoupled, 1P–Coupled, 2P–Uncoupled), averaged across all participants (N = 36, 18 dyads). **Spatial activation patterns** represent the topographical distribution over the scalp of the entrained (top) and beta (bottom) components. For the entrained component, spatial activation patterns were z-score normalized across conditions before averaging, due to the presence of abnormal variance in some participants’ channels. Consistent with the manuscript’s emphasis on temporal dynamics, we refrain from strong claims regarding the anatomical generators of the observed topographies. Nonetheless, activation over the right temporo-parietal junction (TPJ) in the 1P–Coupled condition is noteworthy, as this region has previously been implicated in social interactions, particularly in low-level processing such as the attentional allocation to salient stimuli and attribution of agency ^51^. **Eigenspectra** show the proportion of explained variance for the first 10 components in each condition. For the entrained component, the leading GED eigenvalue clearly dominates, indicating successful separation of the target signal and justifying its use as a spatial filter. In contrast, the eigenspectra for beta components show the eigenvalues of the PCA applied to the top 10 GED beta components to extract the dominant source of variance in beta power. This additional step on beta components was motivated by the gradual decay of GED eigenvalues, suggesting that relevant dynamics were distributed across multiple components which we aggregated with PCA.

### EEG hyperscanning

Following source separation performed on each individual participant, two distinct analyses were performed on the entrained (1.667 Hz or 1.641 Hz) and beta (20 Hz) components within each dyad, for each experimental condition. As compared to traditional hyperscanning methods based on interbrain measures of synchrony ^31^, we advocate for explicitly modelling the dynamics of the ongoing interaction in order to provide meaningfully interpretable measures. *Neural entrainment* within dyads was computed as the convergence of individual instantaneous frequency timeseries ^23,25^ towards a shared frequency. The same Gaussian filter used for GED was applied to the individual entrained components to extract reliable phase timeseries from the analytic signal ^77^, computed via Hilbert transform. In order to remove discontinuities caused by phase resets, the timeseries were unwrapped, differenced, and scaled to Hz ^34^:

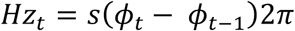

The resulting instantaneous frequency timeseries were smoothed with a sliding moving median (window width of 400 samples), to remove transient artifactual activity that may distort the phase timeseries ^34^.

For each dyad, a time-varying measure of convergence was computed as the difference between the individual instantaneous frequency timeseries: 𝐻𝑧𝑡_Sub#2 - 𝐻𝑧𝑡_Sub#1. A value of 0 was expected in case of perfect convergence, whereas a value of -0.026 Hz was expected in case of maintenance of the assigned metronomes’ frequencies. The resulting convergence timeseries were segmented in 10 trials aligned with each full drifting metronomes’ phase cycle (0 to 2pi), and divided in 64 consecutive phase bins. Individual samples were averaged within each bin. The resulting timeseries shared the same format as the behavioral measures analyzed in our previous works ^13,26,27^, allowing us to replicate the statistical analysis.

*Beta modulation* was computed for every participant based on the distribution of beta power as a function of the phase of the finger-tapping cycles produced by the partner ^24^. In order to compute beta power timeseries, the same plateau-shaped FIR filter used for GED was applied to the beta components before performing the Hilbert transform. For each of the 37 beta components separated by GED, beta power was computed as the squared magnitude of the resulting analytic signal. Given the worse performance of GED for separating beta components as compared to low-frequency entrained components (see Figure 5), principal components analysis (PCA) was performed on the set of power timeseries to extract shared variance of power across all beta components, independently for each participant. The activation timeseries of the 1^st^ principal component (PC#1) was retained, reducing the dimensionality while factoring in the weighted contribution of all beta components. Extreme values deviating from the mean by more than 3 standard deviations were considered outliers and removed from PC#1. Finger-tapping phase timeseries were divided into 36 bins (bin size = 10°) ^42^, so that the median PC#1 values falling within the same bins were computed. With this procedure, we obtained the beta power curves as a function of the partner’s movement cycles, computed over the total amount of finger-taps (645 events were expected on average from each participant). Based on the observation that beta power modulation could be well approximated by a sinusoidal function across one full tapping cycle, the best fitting sinewave was estimated using the *sineFit* Matlab® function ^100^, for each participant in each experimental condition. The sine amplitude provided us with a measure of beta modulation strength, replicating the approach presented in ^24^.

The processing pipelines leading to the computation of neural entrainment and beta modulation are illustrated in Figure 2.

### Statistical modelling

For *neural entrainment*, we used the frequency difference timeseries as the response variable in a mixed-effects model. The model included Coupling and Perspective as factors, and Time as a continuous predictor expressed as the indexes of the metronome steps (from 1 to 64 in one drifting metronomes’ cycle). Due to the non-linearity of the curves, we employed the method of orthogonal polynomials ^50^, incorporating linear and quadratic functions of Time into our model (2^nd^-order polynomial). Dyads, as well as the interactions between Dyads and the factors, were modeled as random effects on all polynomial terms to accommodate individual variability in synchronization abilities and individual susceptibility to coupling across experimental manipulations. The full model’s formula was defined as follows:

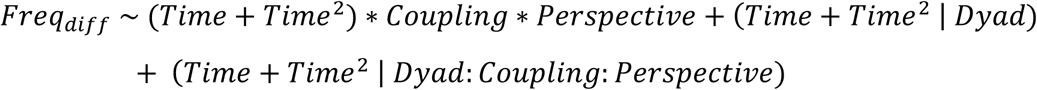

For *beta modulation,* the sine amplitudes quantifying beta modulation strength were log-transformed to ensure residuals met the assumptions of normality (p = 0.057, Shapiro–Wilk test).

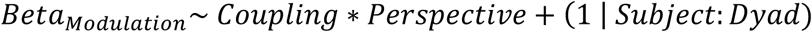

Factors were leveled such that ‘Uncoupled’ and ‘2P’ would provide the baseline intercept for the model, since the modulation by the other’s movements was expected to be null in absence of coupling between the partners. The interaction between individual Subjects and the respective Dyad was modelled as random effect.

EEG processing and analyses were entirely carried out in Matlab® (version R2019a). Statistical analyses were carried out in R (version 4.0.3), using the lme4 package ^101^ for model fitting.

## Data and code availability

The pre-processed EEG data generated and analyzed during this study are available from the corresponding author upon reasonable request.

All analysis scripts are publicly available at: https://github.com/mattiaRosso92/Oscillatory_Dynamics_Joint_Action

## Acknowledgements

The present study was funded by Bijzonder Onderzoeksfonds (BOF) from Ghent University (Belgium), in the context of a joint-PhD project with the University of Lille (France) (l-SITE ULNE program, grant number 01D21819).

The authors are grateful to Canan Nuran Gener for her precious help in collecting the data, to Ivan Schepers for building the hardware of the finger-tapping device, and to Kevin Smink for the illustration of the experimental design.

The Center for Music in the Brain (MIB) is funded by the Danish National Research Foundation (project number DNRF117).

## Author contributions statement

M.R. conceived the hypotheses and designed the study. M.R. and B.V.K. conceived and implemented the experimental setup. M.R. and B.V.K. collected the data. M.R. performed pre- processing and data analysis. M.R. and M.L. performed statistical analysis. P.K., M.L., P.M., and P.V. provided essential help to interpret and frame the results within the neuroscientific literature. M.R. and P.V. wrote the first draft of the manuscript. M.R. prepared the figures. All the authors contributed to and approved the final version of the manuscript.

## Competing interest statement

The authors declare no competing interests.

## Notes

### Competing Interest Statement

The authors have declared no competing interest.

## References

1. Frith, C. D. & Frith, U. Mechanisms of social cognition. Annu. Rev. Psychol. 63, 287–313 (2012).

2. Healey, M. L. & Grossman, M. Cognitive and affective perspective-taking: Evidence for shared and dissociable anatomical substrates. Front. Neurol. 9, 491 (2018).

3. Tversky, B. & Hard, B. M. Embodied and disembodied cognition: spatial perspective-taking. Cognition 110, 124–129 (2009).

4. Kessler, K. & Thomson, L. A. The embodied nature of spatial perspective taking: embodied transformation versus sensorimotor interference. Cognition 114, 72–88 (2010).

5. Furlanetto, T., Bertone, C. & Becchio, C. The bilocated mind: new perspectives on self-localization and self-identification. Front. Hum. Neurosci. 7, 71 (2013).

6. Conson, M. et al. “put myself into your place”: Embodied simulation and perspective taking in autism spectrum disorders: Perspective taking in ASD. Autism Res. 8, 454–466 (2015).

7. Erle, T. M. & Topolinski, S. The grounded nature of psychological perspective-taking. J. Pers. Soc. Psychol. 112, 683–695 (2017).

8. Frith, C. D. & Frith, U. Implicit and explicit processes in social cognition. Neuron 60, 503–510 (2008).

9. Petkova, V. I. & Ehrsson, H. H. If I were you: perceptual illusion of body swapping. PLoS One 3, e3832 (2008).

10. Petkova, V. I., Khoshnevis, M. & Ehrsson, H. H. The perspective matters! Multisensory integration in ego- centric reference frames determines full-body ownership. Front. Psychol. 2, 35 (2011).

11. Petkova, V. I. et al. From part- to whole-body ownership in the multisensory brain. Curr. Biol. 21, 1118– 1122 (2011).

12. Maes, P.-J., van Kerrebroeck, B., Rosso, M., Marouda, I. & Leman, M. Extended reality (XR) in embodied musical art and science. in The Routledge Handbook of Embodied Cognition 143–155 (Routledge, London, 2024).

13. Rosso, M., van Kerrebroeck, B., Maes, P.-J. & Leman, M. Embodied perspective-taking enhances interpersonal synchronization. A body-swap study. iScience (2023).

14. Heggli, O. A., Konvalinka, I., Kringelbach, M. L. & Vuust, P. A metastable attractor model of self–other integration (MEAMSO) in rhythmic synchronization. Philos. Trans. R. Soc. Lond. B Biol. Sci. 376, 20200332 (2021).

15. Heggli, O. A., Cabral, J., Konvalinka, I., Vuust, P. & Kringelbach, M. L. A Kuramoto model of self-other integration across interpersonal synchronization strategies. PLoS Comput. Biol. 15, e1007422 (2019).

16. Heggli, O. A. et al. Transient brain networks underlying interpersonal strategies during synchronized action. Soc. Cogn. Affect. Neurosci. 16, 19–30 (2021).

17. Konvalinka, I., Vuust, P., Roepstorff, A. & Frith, C. D. Follow you, follow me: continuous mutual prediction and adaptation in joint tapping. Q. J. Exp. Psychol. 63, 2220–2230 (2010).

18. Tsakiris, M. My body in the brain: a neurocognitive model of body-ownership. Neuropsychologia 48, 703– 712 (2010).

19. Tsakiris, M. The multisensory basis of the self: From body to identity to others. Q. J. Exp. Psychol. 70, 597– 609 (2017).

20. Mottelson, A., Muresan, A., Hornbæk, K. & Makransky, G. A systematic review and meta-analysis of the effectiveness of body ownership illusions in virtual reality. ACM Trans. Comput. Hum. Interact. (2023) doi:10.1145/3590767.

21. Pyasik, M., Ciorli, T. & Pia, L. Full body illusion and cognition: A systematic review of the literature. Neurosci. Biobehav. Rev. 143, 104926 (2022).

22. Koban, L., Ramamoorthy, A. & Konvalinka, I. Why do we fall into sync with others? Interpersonal synchronization and the brain’s optimization principle. Soc. Neurosci. 14, 1–9 (2019).

23. Rosso, M., Leman, M. & Moumdjian, L. Neural Entrainment Meets Behavior: The Stability Index as a Neural Outcome Measure of Auditory-Motor Coupling. Front. Hum. Neurosci. 15, 668918 (2021).

24. Rosso, M., Heggli, O. A., Maes, P. J., Vuust, P. & Leman, M. Mutual beta power modulation in dyadic entrainment. Neuroimage 257, 119326 (2022).

25. Rosso, M., Moens, B., Leman, M. & Moumdjian, L. Neural entrainment underpins sensorimotor synchronization to dynamic rhythmic stimuli. Neuroimage 277, 120226 (2023).

26. Rosso, M., Maes, P. J. & Leman, M. Modality-specific attractor dynamics in dyadic entrainment. Sci. Rep. 11, 18355 (2021).

27. Rosso, M., Gener, C. N., Moens, B., Maes, P.-J. & Leman, M. Perceptual coupling in human dyads: Kinematics does not affect interpersonal synchronization. Heliyon 10, e33831 (2024).

28. Marsh, K. L., Richardson, M. J. & Schmidt, R. C. Social connection through joint action and interpersonal coordination. Top. Cogn. Sci. 1, 320–339 (2009).

29. Tognoli, E., Zhang, M., Fuchs, A., Beetle, C. & Kelso, J. A. S. Coordination Dynamics: A Foundation for Understanding Social Behavior. Front. Hum. Neurosci. 14, 317 (2020).

30. Tognoli, E. & Kelso, J. A. S. The metastable brain. Neuron 81, 35–48 (2014).

31. Holroyd, C. B. Interbrain synchrony: on wavy ground. Trends Neurosci. 45, 346–357 (2022).

32. Lakatos, P., Gross, J. & Thut, G. A New Unifying Account of the Roles of Neuronal Entrainment. Curr. Biol. 29, R890–R905 (2019).

33. Haegens, S. & Zion Golumbic, E. Rhythmic facilitation of sensory processing: A critical review. Neurosci. Biobehav. Rev. 86, 150–165 (2018).

34. Cohen, M. X. Fluctuations in oscillation frequency control spike timing and coordinate neural networks. J. Neurosci. 34, 8988–8998 (2014).

35. Miyata, K., Varlet, M., Miura, A., Kudo, K. & Keller, P. E. Modulation of individual auditory-motor coordination dynamics through interpersonal visual coupling. Sci. Rep. 7, 16220 (2017).

36. Miyata, K., Varlet, M., Miura, A., Kudo, K. & Keller, P. E. Interpersonal visual interaction induces local and global stabilisation of rhythmic coordination. Neurosci. Lett. 682, 132–136 (2018).

37. Bigand, F., Bianco, R., Abalde, S. F. & Novembre, G. The geometry of interpersonal synchrony in human dance. Curr. Biol. 34, 3011–3019.e4 (2024).

38. Seeber, M., Scherer, R., Wagner, J., Solis-Escalante, T. & Müller-Putz, G. R. EEG beta suppression and low gamma modulation are different elements of human upright walking. Front. Hum. Neurosci. 8, 485 (2014).

39. Seeber, M., Scherer, R. & Müller-Putz, G. R. EEG oscillations are modulated in different behavior-related networks during rhythmic finger movements. Journal of Neuroscience (2016).

40. Fujioka, T., Trainor, L., Large, E. & Ross, B. Beta and gamma rhythms in human auditory cortex during musical beat processing. Ann. N. Y. Acad. Sci. (2009).

41. Fujioka, T., Trainor, L. J., Large, E. W. & Ross, B. Internalized Timing of Isochronous Sounds Is Represented in Neuromagnetic Beta Oscillations. J. Neurosci. 32, 1791–1802 (2012).

42. Zhou, G., Bourguignon, M., Parkkonen, L. & Hari, R. Neural signatures of hand kinematics in leaders vs. followers: A dual-MEG study. Neuroimage 125, 731–738 (2016).

43. van Ede, F., Szebényi, S. & Maris, E. Attentional modulations of somatosensory alpha, beta and gamma oscillations dissociate between anticipation and stimulus processing. Neuroimage 97, 134–141 (2014).

44. Novembre, G., Knoblich, G., Dunne, L. & Keller, P. E. Interpersonal synchrony enhanced through 20 Hz phase-coupled dual brain stimulation. Soc. Cogn. Affect. Neurosci. (2017) doi:10.1093/scan/nsw172.

45. Abalde, S. F., Rigby, A., Keller, P. E. & Novembre, G. A framework for joint music making: Behavioral findings, neural processes, and computational models. Neurosci. Biobehav. Rev. 167, 105816 (2024).

46. Zamm, A., Debener, S., Konvalinka, I., Sebanz, N. & Knoblich, G. The sound of silence: an EEG study of how musicians time pauses in individual and joint music performance. Soc. Cogn. Affect. Neurosci. 16, 31– 42 (2021).

47. Ménoret, M., Bourguignon, M. & Hari, R. Modulation of rolandic beta-band oscillations during motor simulation of joint actions. PLoS One 10, e0131655 (2015).

48. Krol, M. A. & Jellema, T. Sensorimotor representation of observed dyadic actions with varying agent involvement: an EEG mu study. Cogn. Neurosci. 14, 25–35 (2023).

49. Bolt, N. K. & Loehr, J. D. Motor-related cortical oscillations distinguish one’s own from a partner’s contributions to a joint action. Biol. Psychol. 190, 108804 (2024).

50. Mirman, D. Growth Curve Analysis and Visualization Using R. (CRC Press, 2017).

51. Decety, J. & Lamm, C. The role of the right temporoparietal junction in social interaction: how low-level computational processes contribute to meta-cognition. Neuroscientist 13, 580–593 (2007).

52. Iversen, J. R., Repp, B. H. & Patel, A. D. Top-down control of rhythm perception modulates early auditory responses. Ann. N. Y. Acad. Sci. 1169, 58–73 (2009).

53. Criscuolo, A., Schwartze, M., Henry, M. J., Obermeier, C. & Kotz, S. A. Individual neurophysiological signatures of spontaneous rhythm processing. Neuroimage 273, 120090 (2023).

54. Oouchida, Y. et al. Maladaptive change of body representation in the brain after damage to central or peripheral nervous system. 104, 38–43 (2016).

55. Maselli, A. & Slater, M. The building blocks of the full body ownership illusion. Front. Hum. Neurosci. 7, 83 (2013).

56. Slater, M., Spanlang, B., Sanchez-Vives, M. V. & Blanke, O. First person experience of body transfer in virtual reality. PLoS One 5, e10564 (2010).

57. Samad, M., Chung, A. J. & Shams, L. Perception of body ownership is driven by Bayesian sensory inference. PLoS One 10, e0117178 (2015).

58. Dupraz, L., Bourgin, J., Pia, L., Barra, J. & Guerraz, M. Body ownership and kinaesthetic illusions: Dissociated bodily experiences for distinct levels of body consciousness? Conscious. Cogn. 117, 103630 (2024).

59. Kalckert, A. & Ehrsson, H. H. Moving a Rubber Hand that Feels Like Your Own: A Dissociation of Ownership and Agency. Front. Hum. Neurosci. 6, 40 (2012).

60. Kalckert, A. & Ehrsson, H. H. The moving rubber hand illusion revisited: comparing movements and visuotactile stimulation to induce illusory ownership. Conscious. Cogn. 26, 117–132 (2014).

61. Harry, B. & Keller, P. E. Tutorial and simulations with ADAM: an adaptation and anticipation model of sensorimotor synchronization. Biol. Cybern. 113, 397–421 (2019).

62. Novembre, G., Sammler, D. & Keller, P. E. Neural alpha oscillations index the balance between self-other integration and segregation in real-time joint action. Neuropsychologia 89, 414–425 (2016).

63. Keitel, A. & Gross, J. Individual Human Brain Areas Can Be Identified from Their Characteristic Spectral Activation Fingerprints. PLoS Biol. 14, e1002498 (2016).

64. Rosso, M. et al. FREQ-NESS reveals the dynamic reconfiguration of frequency-resolved brain networks during auditory stimulation. Adv. Sci. (Weinh.) e2413195 (2025).

65. Bourguignon, M., Jousmäki, V., Dalal, S. S., Jerbi, K. & De Tiège, X. Coupling between human brain activity and body movements: Insights from non-invasive electromagnetic recordings. Neuroimage 203, 116177 (2019).

66. Baker, M. R. & Baker, S. N. The effect of diazepam on motor cortical oscillations and corticomuscular coherence studied in man. J. Physiol. 546, 931–942 (2003).

67. Baker, S. N., Chiu, M. & Fetz, E. E. Afferent encoding of central oscillations in the monkey arm. J. Neurophysiol. 95, 3904–3910 (2006).

68. Baker, S. N. Oscillatory interactions between sensorimotor cortex and the periphery. 17, 649–655 (2007).

69. Witham, C. L., Riddle, C. N., Baker, M. R. & Baker, S. N. Contributions of descending and ascending pathways to corticomuscular coherence in humans: Descending and ascending corticomuscular coherence. J. Physiol. 589, 3789–3800 (2011).

70. Novembre, G. & Iannetti, G. D. Hyperscanning Alone Cannot Prove Causality. Multibrain Stimulation Can. Trends Cogn. Sci. 25, 96–99 (2021).

71. Pfurtscheller, G. Central beta rhythm during sensorimotor activities in man. Electroencephalogr. Clin. Neurophysiol. 51, 253–264 (1981).

72. Toma, K. et al. Movement rate effect on activation and functional coupling of motor cortical areas. J. Neurophysiol. 88, 3377–3385 (2002).

73. Morillon, B., Arnal, L. H., Schroeder, C. E. & Keitel, A. Prominence of delta oscillatory rhythms in the motor cortex and their relevance for auditory and speech perception. Neurosci. Biobehav. Rev. 107, 136–142 (2019).

74. MacDougall, H. G. & Moore, S. T. Marching to the beat of the same drummer: the spontaneous tempo of human locomotion. J. Appl. Physiol. 99, 1164–1173 (2005).

75. Phillips-Silver, J., Aktipis, C. A. & Bryant, G. A. The ecology of entrainment: Foundations of coordinated rhythmic movement. Music Percept. 28, 3–14 (2010).

76. Rajendran, V. G. & Schnupp, J. W. H. Frequency tagging cannot measure neural tracking of beat or meter. Proceedings of the National Academy of Sciences of the United States of America vol. 116 2779–2780 (2019).

77. Rosenblum, M., Pikovsky, A., Kurths, J., Schäfer, C. & Tass, P. A. Chapter 9 Phase synchronization: From theory to data analysis. in Handbook of Biological Physics (eds. Moss, F. & Gielen, S.) vol. 4 279–321 (North-Holland, 2001).

78. Richardson, M. J., Marsh, K. L. & Baron, R. M. Judging and actualizing intrapersonal and interpersonal affordances. J. Exp. Psychol. Hum. Percept. Perform. 33, 845–859 (2007).

79. Zalta, A., Petkoski, S. & Morillon, B. Natural rhythms of periodic temporal attention. Nat. Commun. 11, 1051 (2020).

80. Zalta, A., Large, E. W., Schön, D. & Morillon, B. Neural dynamics of predictive timing and motor engagement in music listening. Sci. Adv. 10, eadi2525 (2024).

81. Large, E. W. & Jones, M. R. The dynamics of attending: How people track time-varying events. Psychol. Rev. 106, 119–159 (1999).

82. Henry, M. & Herrmann, B. Low-frequency neural oscillations support dynamic attending in temporal context. Timing & Time Perception 2, 62–86 (2014).

83. Lakatos, P. et al. An oscillatory hierarchy controlling neuronal excitability and stimulus processing in the auditory cortex. J. Neurophysiol. 94, 1904–1911 (2005).

84. Schroeder, C. E. & Lakatos, P. Low-frequency neuronal oscillations as instruments of sensory selection. Trends Neurosci. 32, 9–18 (2009).

85. Lakatos, P., Karmos, G., Mehta, A. D., Ulbert, I. & Schroeder, C. E. Entrainment of neuronal oscillations as a mechanism of attentional selection. Science 320, 110–113 (2008).

86. Busch, N., Dubois, J. & VanRullen, R. The phase of ongoing EEG oscillations predicts visual perception. J. Neurosci. 29, 7869–7876 (2009).

87. Busch, N. A. & VanRullen, R. Spontaneous EEG oscillations reveal periodic sampling of visual attention. Proc. Natl. Acad. Sci. U. S. A. 107, 16048–16053 (2010).

88. Vanrullen, R., Busch, N. A., Drewes, J. & Dubois, J. Ongoing EEG phase as a trial-by-trial predictor of perceptual and attentional variability. Front. Psychol. 2, 60 (2011).

89. Henry, M. J. & Obleser, J. Frequency modulation entrains slow neural oscillations and optimizes human listening behavior. Proc. Natl. Acad. Sci. U. S. A. 109, 20095–20100 (2012).

90. Morillon, B. & Baillet, S. Motor origin of temporal predictions in auditory attention. Proc. Natl. Acad. Sci. U. S. A. 114, E8913–E8921 (2017).

91. Shinozuka, K., et al. LSD reconfigures the frequency-specific network landscape of the human brain. bioRxiv (2025) doi:10.1101/2025.03.21.644645.

92. Bonetti, L. et al. BROADband brain Network Estimation via Source Separation (BROAD-NESS). bioRxiv (2024) doi:10.1101/2024.10.31.621257.

93. Cohen, M. X. & Gulbinaite, R. Rhythmic entrainment source separation: Optimizing analyses of neural responses to rhythmic sensory stimulation. Neuroimage 147, 43–56 (2017).

94. Cohen, M. X. A tutorial on generalized eigendecomposition for denoising, contrast enhancement, and dimension reduction in multichannel electrophysiology. Neuroimage 247, 118809 (2022).

95. Oostenveld, R., Fries, P., Maris, E. & Schoffelen, J.-M. FieldTrip: Open source software for advanced analysis of MEG, EEG, and invasive electrophysiological data. Comput. Intell. Neurosci. 2011, 156869 (2011).

96. Makeig, S., Bell, A. J., Jung, T. & Sejnowski, T. Independent Component Analysis of electroencephalographic data. Neural Inf Process Syst 8, 145–151 (1995).

97. Makeig, S., Jung, T. P., Bell, A. J., Ghahremani, D. & Sejnowski, T. J. Blind separation of auditory event-related brain responses into independent components. Proc. Natl. Acad. Sci. U. S. A. 94, 10979–10984 (1997).

98. Vidal, M., Rosso, M. & Aguilera, A. M. Bi-smoothed functional independent component analysis for eeg artifact removal. Sci. China Ser. A Math. (2021).

99. Cohen, M. X. A data-driven method to identify frequency boundaries in multichannel electrophysiology data. J. Neurosci. Methods 347, 108949 (2021).

100. Seibold, P. Sine Fitting (https://Www.Mathworks.Com/Matlabcentral/Fileexchange/66793-Sine-Fitting), MATLAB Central File Exchange. (2021).

101. Bates, D., Mächler, M., Bolker, B. & Walker, S. Fitting Linear Mixed-Effects Models using lme4. arXiv [stat.CO] (2014).

